# Phylogenomic comparative methods: accurate evolutionary inferences in the presence of gene tree discordance

**DOI:** 10.1101/2022.11.14.516436

**Authors:** Mark S. Hibbins, Lara C. Breithaupt, Matthew W. Hahn

## Abstract

Phylogenetic comparative methods have long been a mainstay of evolutionary biology, allowing for inferences of the tempo and mode of trait evolution across species while accounting for their common ancestry. These analyses typically assume a single, bifurcating phylogenetic tree that describes the shared history among species. However, modern phylogenomic analyses have shown that genomes are often composed of a mosaic of different histories that can disagree both with the species tree and with each other—so-called discordant gene trees. These gene trees describe shared histories that are not captured by the species tree, and therefore that are unaccounted for in classic comparative approaches. The application of standard phylogenetic comparative methods to species histories containing discordance leads to incorrect inferences about the timing, direction, and rate of evolution. Here, we develop two approaches for incorporating gene tree histories into comparative methods: one involves constructing a fuller phylogenetic variance-covariance matrix that includes relationships not found in the species tree, and another that applies Felsenstein’s pruning algorithm over a set of gene trees to calculate trait histories and likelihoods. Both approaches are agnostic to the biological causes of gene tree discordance, which may include incomplete lineage sorting and introgression. Using simulation, we demonstrate that our new approaches generate much more accurate estimates of tree-wide rates of trait evolution than standard methods. We apply our methods to two clades of the wild tomato genus *Solanum* with varying rates of discordance, demonstrating the contribution of gene tree discordance to variation in a set of floral traits and the ability of our approaches to provide more accurate inferences. Our new approaches have the potential to be applied to a broad range of classic inference problems in phylogenetics, including ancestral state reconstruction and the inference of lineage-specific rate shifts.

## Introduction

A major goal of evolutionary biology is to understand how and why traits vary among species. One of the major sources of this variation is common ancestry. If left unaccounted for, this shared history can lead to pseudoreplication and spurious trait correlations (Felsenstein 1985). Phylogenetic comparative methods have been developed to account for shared history, enabling more accurate inferences about the tempo and mode of trait evolution (Harvey and Pagel 1991). With the statistical toolkit offered by phylogenetic comparative methods, researchers can ask questions about the rate at which traits evolve, whether these rates have changed over time or in different lineages, what traits may have looked like in ancestral or extinct lineages, and whether trait shifts are correlated with historical or environmental factors (Martins and Hansen 1996; Garamszegi 2014; Adams and Collyer 2018; Revell and Harmon 2022).

In classic comparative methods, common ancestry among species is accounted for using a single species phylogeny. However, genome-scale analyses of phylogenetic history have revealed that individual loci can have their own independent histories (Pollard et al. 2006; White et al. 2009; Fontaine et al. 2015; Pease et al. 2016; Novikova et al. 2016; Copetti et al. 2017; Wu et al. 2018; Edelman et al. 2019; Vanderpool et al. 2020). The result is gene tree discordance—the disagreement of trees at individual loci both with each other and with the species phylogeny. Gene tree discordance has important implications for phylogenetic comparative methods because discordant gene trees contain branches that are not present in the species phylogeny. Evolution along such discordant branches can result in trait similarity among species with no shared history in the species tree (Figure 1). Such patterns of trait variation can mislead standard phylogenetic comparative methods, particularly by resulting in overestimates of the number of trait transitions or the rate of trait evolution (Hahn and Nakhleh 2016; Mendes and Hahn 2016; Mendes et al. 2018; Hibbins et al. 2020; Wang et al. 2021). This effect has been termed “hemiplasy,” as single transitions on discordant gene trees can falsely resemble homoplasy when analyzed on the species tree (Avise and Robinson 2008).

**Figure 1:**
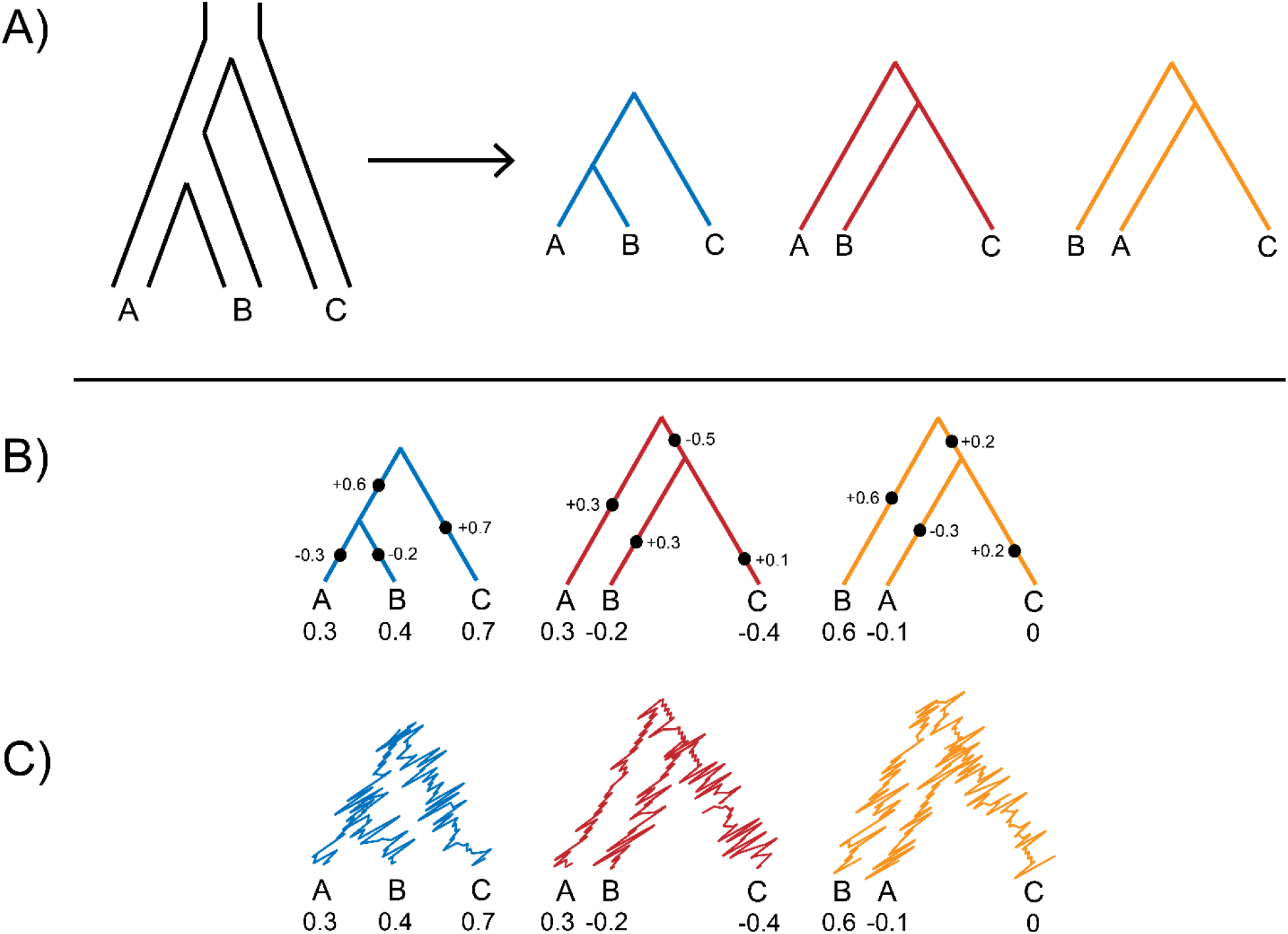
Conceptualizing quantitative trait evolution with discordant gene trees. A) Given a species tree (far left), we model gene trees as arising under the multispecies coalescent process. One topology is concordant with the species tree (blue), while the other two possible topologies are discordant (red and yellow). Under ILS these two discordant trees have the same topology and frequency. B) Over the course of evolution, mutations occur at loci that affect quantitative traits, each of which has a topology drawn from the multispecies coalescent process. Mutations on the internal branches of discordant gene trees can introduce shared trait history that is not captured by the species tree. Here we summarize mutations occurring at different loci on a single tree if the loci had the same topology, with each mutation at each locus contributing positively or negatively to the trait value in each species. In the example here, species pairs B-C and A-C might covary in quantitative trait values, despite sharing no common ancestor in the species tree. C) Given a large number of mutations and loci, trait evolution over time can be modelled by Brownian motion on each gene tree topology. This stochastic process models the trait value as a random walk over time, with species trait values calculated as the weighted average of the values on each gene tree (Mendes et al. 2018; Hibbins and Hahn 2021).

Discordance is a concern for evolutionary inference because it has biological causes that cannot be overcome by addressing technical errors or by increasing species sampling (Degnan and Rosenberg 2009). Two primary causes of discordance, incomplete lineage sorting (ILS) and introgression, have different effects on gene tree frequencies and branch lengths and are therefore expected to bias comparative methods in different ways. ILS, a stochastic process that depends on species tree internal branch lengths and population sizes, generates symmetry in the frequencies of possible discordant gene trees (Hudson 1983; Pamilo and Nei 1988). Therefore, higher amounts of ILS lead to broad increases in the occurrence of hemiplasy across multiple possible incongruent trait patterns (Guerrero and Hahn 2018; Mendes et al. 2018). Introgression is a process of historical hybridization and back-crossing that, while widespread in modern phylogenomic datasets (Mallet et al. 2016; Taylor and Larson 2019; Dagilis et al. 2022), is often more limited to specific pairs of taxa. In particular, post-speciation introgression between non-sister lineages leads to an excess of gene trees grouping those lineages as sister (Reich et al. 2009; Green et al. 2010; Durand et al. 2011; Patterson et al. 2012). This pattern should result in an excess of trait-sharing for the species involved in introgression compared to the species not exchanging genes (Hibbins et al. 2020; Hibbins and Hahn 2021; Wang et al. 2021).

While some progress has been made in accounting for discordance in the evolution of discrete traits, especially in nucleotide models (De Maio et al. 2013; De Maio et al. 2015; Schrempf et al. 2016; Ogilvie et al. 2017; Schrempf et al. 2019), many classic phylogenetic comparative methods remain unable to account for gene tree discordance when analyzing quantitative traits. The approaches required to improve these methods will depend on the question being asked. Some tasks, such as maximum-likelihood estimation of the rate of trait evolution under Brownian motion (σ^2^) (e.g. Garland and Ives 2000; O’Meara et al. 2006) or phylogenetic regression (Grafen 1989), depend on the specification of a matrix that describes the trait variances and covariances expected from the species phylogeny (often denoted ***C***). Other comparative approaches, such as ancestral state reconstruction (Pagel 1999) and inference of lineage-specific rate shifts (Alfaro et al. 2009), can require more sophisticated approaches that calculate state probabilities on different parts of a phylogeny; one such approach is to use Felsenstein’s pruning algorithm applied to a species tree with specified branch lengths (Felsenstein 1973). Mendes et al. (2018) showed that failing to account for discordance can bias estimates of σ^2^ upwards and can lead to falsely inflated numbers of trait-mean transitions. In general, the development of a comparative framework incorporating gene tree discordance would lead to more accurate evolutionary inferences in a wide variety of systems with ILS and/or introgression, across a wide variety of approaches for making inferences about quantitative traits.

Here, we demonstrate the utility of an updated *phylogenomic* comparative framework, using two distinct approaches to incorporate the summed history of concordant and discordant gene trees into evolutionary inference. In the first approach, we show how to construct an updated phylogenetic variance/covariance matrix (which we denote ***C*\***) to include the covariances introduced by discordant gene trees. We provide a new R package, *seastaR*, that can construct this updated matrix for any number of species, either by summing the internal branches of an input set of gene trees or by calculating expected gene tree internal branches from an input species tree using the multispecies coalescent model. We show how estimates of the evolutionary rate are made more accurate by using ***C*\***, and suggest how this updated matrix can be passed to other available software packages to make multiple evolutionary inferences more robust to discordance. In the second approach, we develop a method for applying the pruning algorithm over a set of gene trees to return the likelihood of an observed trait across species. Using a pilot implementation of this approach for a rooted three-species tree, we show how it can be used to accurately estimate the rate of quantitative trait evolution. Although currently limited to a smaller number of species, this latter approach has the potential to perform more complicated comparative inferences in the presence of discordance. Finally, we apply our approaches to empirical morphological data from wild tomatoes (Haak et al. 2014), finding a greater discrepancy between species tree and gene tree rate estimates in a clade with a higher rate of gene tree discordance. Overall, our new approaches pave the way towards more accurate evolutionary inferences in the presence of gene tree discordance.

## Methods

### Building a phylogenetic variance/covariance matrix from data with discordance

As previously discussed, one of the most common ways that phylogeny is incorporated into comparative analyses is by constructing a phylogenetic variance/covariance matrix, ***C***. This square matrix has rows and columns corresponding to the number of taxa in the phylogeny, with the diagonal elements containing the expected trait variances for each species and the off-diagonal elements containing the expected trait covariances between each species pair. Considering three species with the relationship ((A,B),C) (Figure 2A), the standard covariance matrix has the following form:

**Figure 2:**
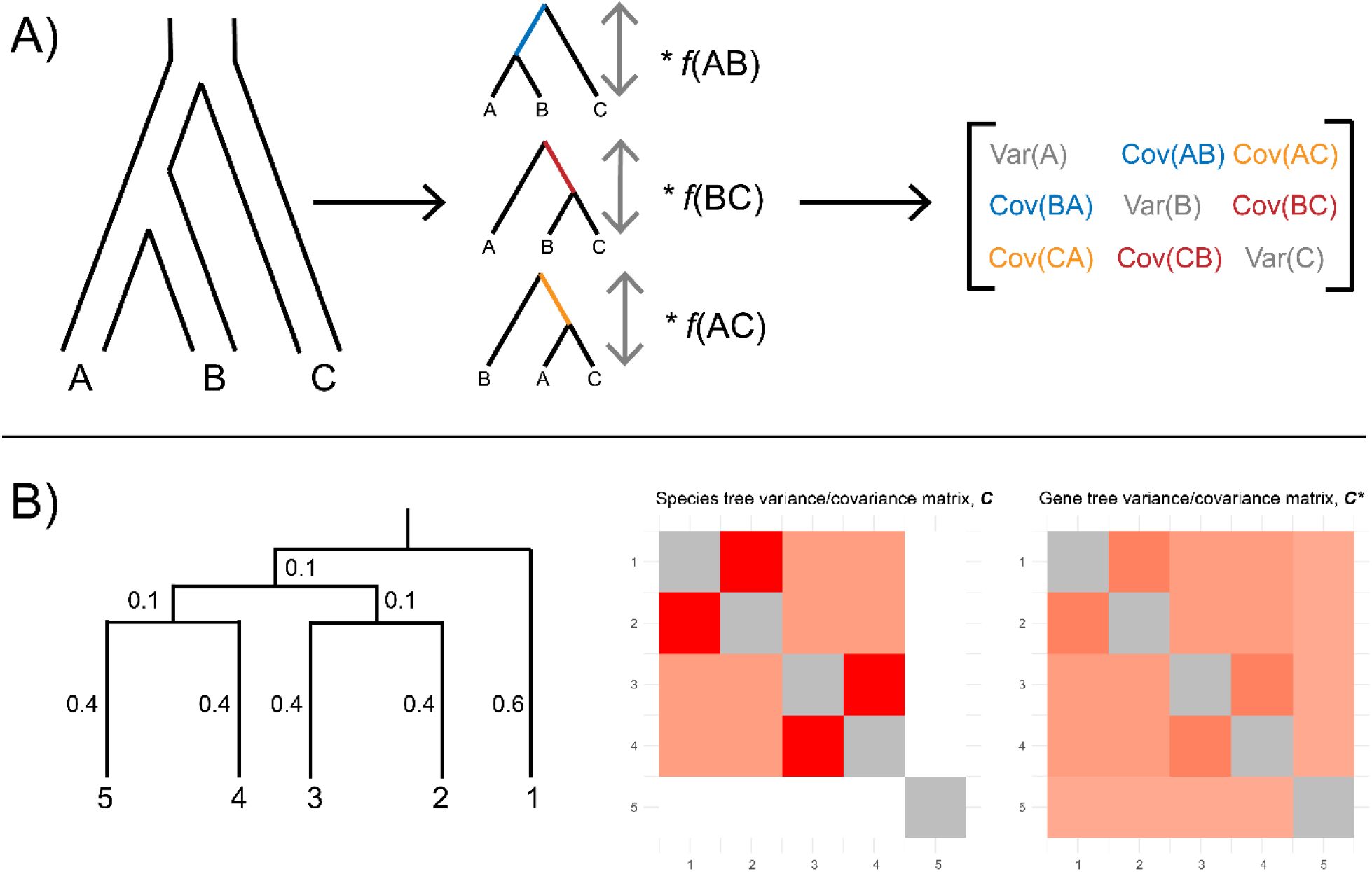
Inferring the gene tree variance/covariance matrix, ***C\****. A) Gene trees are generated from a species tree under the multispecies coalescent process (note that introgression can readily be incorporated, but is not shown here for clarity). Each gene tree contributes its internal branch length (for covariance terms) and its total height (for variance terms) to ***C\****. The contribution of each tree to ***C\**** is weighted by its expected or observed frequency, depending on the approach taken. Frequencies are denoted as *f*(XY), where X and Y are the taxa sister in the gene tree of interest. B) A comparison of ***C*** and ***C\**** for a five-taxon species tree with branch lengths labelled in coalescent units. Each internal branch has a length of 0.1, corresponding to a level of discordance of approximately 60%. This level of discordance means that each clade descended from these internal branches (5/4/3/2, 5/4, and 3/2) will be present in ∼40% of gene trees. The standard phylogenetic covariance matrix, ***C***, contains no covariance between species 1 and the other taxa, because they do not share an internal branch in the species tree. In contrast, species 1 covaries with all other species in the tree using ***C\****, because multiple discordant gene trees have species 2-5 sharing an internal branch with species 1. ***C\**** also has lower covariance between the taxa that are sister in the species tree (5/4+3/2, 5/4, and 3/2).

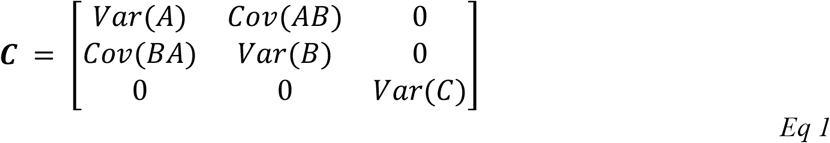

Trait covariances arise from shared internal branches in the phylogeny. As only species A and B share an internal branch in the species tree, the other two species pairs have no expected covariance.

In contrast, if we consider the gene trees that are generated by the species tree in Figure 2A, the two discordant gene trees contain internal branches shared by pairs B-C and A-C. Discordance due to ILS generates all three possible topologies for this species tree, so all off-diagonal entries in the covariance matrix should have non-zero values (Mendes et al. 2018). We are interested in estimating this updated covariance matrix, which we denote ***C\**:**

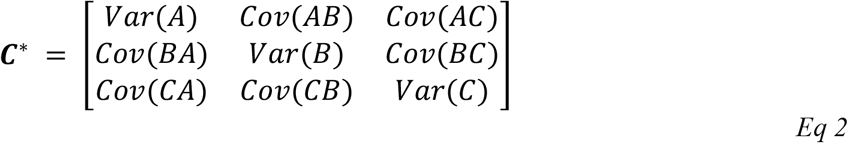

To construct ***C\****, we provide the R package *seastaR. seastaR* uses two approaches for estimating ***C\****, both following the same principle: each gene tree topology contributes an internal branch which, after being weighted by that tree’s expected frequency, fills an off-diagonal entry in the covariance matrix (Figure 2A). Each gene tree also contributes its total height, weighted by frequency, to the expected trait variances for each species.

The first approach for estimating ***C\****, *trees_to_vcv*, constructs this matrix from a list of provided gene trees (with branch lengths) and their observed frequencies. The method works by obtaining all the internal branch lengths present in each gene tree, as well as the height of each gene tree, and averaging them to get ***C\****. A major advantage of this approach is that it can easily account for both ILS and introgression as sources of gene tree discordance, as the effects of both are captured in the distribution of observed gene tree topologies and branch lengths. On the other hand, individual gene trees may be inferred with error, making their branch lengths and frequencies less reliable. If accurately estimated gene trees are unavailable, our second approach, *get_full_matrix*, constructs ***C\**** solely from an input species tree in coalescent units. This method breaks the input phylogeny down into each possible triplet, and for each triplet uses expectations from the multispecies coalescent model to calculate the expected internal branches and frequencies for each possible gene tree (see Mendes et al. 2018). For an exemplar five-taxon tree specified in coalescent units, we compared the standard ***C*** matrix to a ***C\**** matrix computed using *get_full_matrix* (Figure 2B). The test tree has three internal branches, each of length of 0.1 coalescent units. Given these branch lengths, we expect 60% of trees to be discordant for each of these three branches, meaning that only 40% of gene trees will have (for instance) the clade containing species 5 and 4 sister to the clade containing species 3 and 2. As expected, ***C\**** contains covariance entries for species pairs that do not share an internal branch in the species tree, but that share internal branches in at least one discordant gene tree (Figure 2B). In addition, the sister lineages in the species tree have smaller covariances in ***C\**** than in ***C***, because they do not share an internal branch in many discordant trees.

Our package, *seastaR*, contains several other utilities, including a parser for an input set of estimated gene trees, a simulator that can simulate trait evolution using ***C\****, and a function to obtain the maximum-likelihood estimate of σ^2^ using ***C\**** (see Results). Also note that, although not currently implemented, *seastaR* could be extended to construct ***C\**** from an input species network specified in coalescent units, using expectations from the multispecies network coalescent model (Hibbins and Hahn 2021).

### Calculating trait likelihoods over a set of gene trees using Felsenstein’s pruning algorithm

Updating the phylogenetic variance/covariance matrix provides a straightforward solution to accounting for gene tree discordance that works for several important inference tasks in comparative methods. However, many questions require more sophisticated models that do not have straightforward solutions making use of this matrix. For these questions, the field would benefit from a general approach to calculating likelihoods given a set of gene trees and a model of trait evolution. Our solution makes use of Felsenstein’s pruning algorithm (Felsenstein 1973), a dynamic programming algorithm that calculates probabilities for a set of character states across all nodes in a phylogeny. A tree-wide likelihood can be calculated from the probabilities at the root, which can be used in conjunction with numerical optimization methods to estimate model parameters.

We developed an approach to apply the pruning algorithm to a specified set of gene trees, rather than to a single tree. This approach is implemented in C++ and draws heavily on the infrastructure of *CAFE* (Hahn et al. 2005, Mendes et al. 2020), a program that uses the pruning algorithm to calculate likelihoods for a birth-death model of gene family evolution. We make several modifications based on the methods presented in Boucher and Démery (2016) and implemented in *CAGEE* (https://github.com/hahnlab/CAGEE) that allow *CAFE*’s implementation of the pruning algorithm to be applied to continuous traits rather than integer counts of gene families (see also Bertram et al. 2022). First, the pruning algorithm requires a vector of possible discrete character states over which probabilities can be calculated. To obtain this vector from an observed continuous trait, we take the range (−2(*max*(|***X***|), 2(*max*(|***X***|)) where ***X*** is the vector of observed characters for each species. The vector of character states is then filled with 100 equidistant steps from the lower bound to the higher bound. Second, we need to assign probabilities to all the character states at the tips of the phylogeny, so that the pruning algorithm has a place to start. This is straightforward for integer count data, as the observed value can simply be assigned a probability of 1 at the tip. However, for continuous traits it will often be the case that none of the values in our discretized trait vector exactly match the observed values at the tips. Therefore, we implement an approach that distributes the probability at the tip over the two states closest to each of the observed values, proportional to how distant they are relative to each other (Equation 18 in the appendix of Boucher and Démery 2016). Third, to calculate the transition probability between each pair of discretized trait values over a branch in the phylogeny, we use the Brownian motion model. The probability density for Brownian motion is:

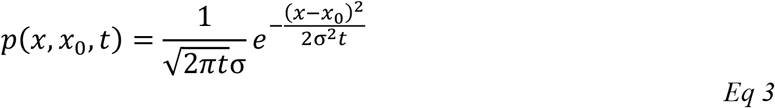

where *x*_0_ is the initial trait value, *x* is the trait value after time *t*, and σ^2^ is the evolutionary rate per unit time. With these methods in place, we can apply the standard pruning algorithm to an individual tree with observed character states and a specified σ^2^ value.

To estimate a single likelihood over a set of gene trees, we initially apply the standard pruning algorithm to each gene tree individually. These gene trees with branch lengths are given to the method directly, and must be ultrametric. Any set of trees can be specified, but the manner in which they are specified will depend on the size of the species tree (i.e. number of tips). For a large species tree, individual gene trees can be inferred or predicted by theory, similarly to the two approaches used by *seastaR*. Because it may not be possible to sample every possible topology, we recommend sampling a reasonable number of individual gene trees (see Discussion). For a small species tree, the most efficient approach will be to specify one tree for each possible topology, along with its frequency. Again, the branch lengths and frequencies of each tree topology can be averaged from a set of inferred trees or predicted from theory. The total likelihood is then calculated as:

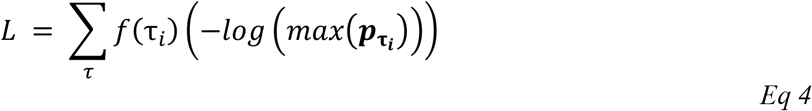

where τ is the set of gene trees, *f*(τ_*i*_) is the frequency of gene tree *i*, and 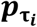 is the vector of character state probabilities at the root for gene tree *i*. In words: we obtain a partial negative log-likelihood for each individual gene tree, these partial likelihoods are then weighted by each gene tree’s observed frequency, and finally the weighted partial likelihoods are summed together to produce the total likelihood (Figure 3).

**Figure 3:**
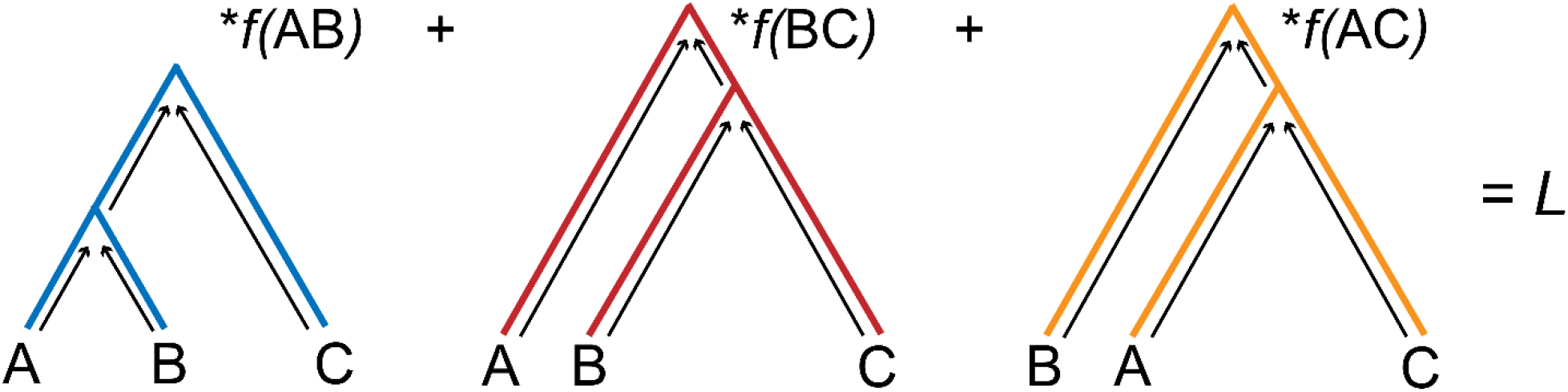
Applying the pruning algorithm to sets of gene trees. In our proposed approach, the pruning algorithm (shown as upward arrows) is applied to each individual gene tree to obtain probabilities at the root for a quantitative trait. These root probabilities are then used to obtain a partial likelihood from each gene tree, which are then summed together weighted by the gene tree frequency to obtain the final likelihood.

A major advantage of the pruning algorithm method is that maximum-likelihood inference can be used to estimate parameters for a wide variety of models. In addition, like the *trees_to_vcv* method of *seastaR*, this approach can easily handle introgression events if the signals of introgression are contained in the specified gene trees, or if expected gene trees under the multispecies network coalescent could be specified by the user. Currently, our implementation uses the Nelder-Mead algorithm (Nelder and Mead 1965) to find the optimal the Brownian motion evolutionary rate parameter, σ^2^. In the future, we would like the software to also perform more sophisticated inferences, such as ancestral state reconstruction or lineage-specific rate shifts. Our approach could also be extended to any evolutionary model, not just Brownian motion.

### Simulating complex traits with discordance

To demonstrate the utility of our new phylogenomic comparative approaches, we used simulations to evaluate their performance on a simple inference task: estimating the evolutionary rate parameter, σ^2^. We simulated traits from a phylogenetic history with increasing rates of gene tree discordance by making random draws from a multivariate normal distribution (where ***C\**** specifies the covariance structure). This simulation approach assumes an infinitesimal contribution to the trait from all genomic loci, an approximation that holds reasonably well for many complex quantitative traits. For each simulated dataset, we applied both standard inference of σ^2^ using the species tree and our updated inferences that account for gene tree discordance. See the Supplementary Methods for the exact conditions and parameters used in our simulations.

We simulated our traits under the model parameterization of Mendes et al. (2018). Levels of discordance in this model are altered by changing the effective population size, *N*, allowing us to increase the level of discordance by increasing *N*. The equations for trait variances and covariances are also scaled by *N*, such that branch lengths in units of absolute time are divided by 2*N*. The evolutionary rate is a compound parameter, 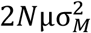, where *μ* is the mutation rate and 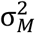 is the variance in mutational effect sizes. A consequence of this formulation of the evolutionary rate is that the true rate used to simulate the data increases as we increase the rate of discordance by increasing *N*. This model is akin to the one shown in Figure 1B, in which mutations occur on gene tree branches with normally distributed effect sizes. Given enough mutations and enough time, these cumulative effects resemble Brownian motion of trait means along each lineage (Figure 1C; Mendes et al. 2018).

### Data availability

Source code and analysis scripts related to *seastaR* and the covariance matrix method can be found in https://github.com/larabreithaupt/seastaR. Code and scripts related to the pruning algorithm method can be found in https://github.com/mhibbins/genetreepruningalg.

## Results

### Phylogenomic comparative approaches yield more accurate evolutionary rate estimates in the presence of discordance

We applied both of our new phylogenomic comparative approaches to data simulated with discordance in order to evaluate their accuracy in estimating the evolutionary rate parameter, σ^2^. For the approach that uses the updated the variance/covariance matrix, ***C\****, we use the maximum-likelihood estimator of σ^2^:

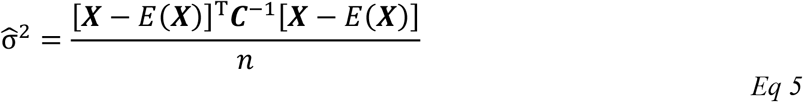

(O’Meara et al. 2006), where ***X*** is the vector of observed trait values at the tips, *C* is the phylogenetic variance-covariance matrix, and *n* is the number of tips. *i****X****i* is the vector containing the expected trait value at the root, calculated as follows:

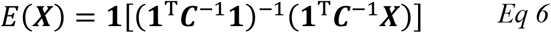

Where **1** is a column vector of ones of size *n* x 1. To account for gene tree discordance with this estimator, we simply use ***C\**** in place of ***C*** in equations 5 and 6. We have implemented this method in *seastaR* to allow users to estimate σ^2^. For this approach, we simulated 1000 trait datasets for each condition of increasing gene tree discordance, estimating σ^2^ using both ***C*** and ***C\**** for each dataset.

For the approach using the pruning algorithm, we implemented the Nelder-Mead optimization algorithm. Given a set of input gene trees and tree frequencies, our optimization approach proposes a new value of σ^2^ in each iteration, returning a single likelihood value over the set of gene trees each time; the optimal value of σ^2^ is the one that maximizes this total likelihood. Owing to longer computation times, we simulated 100 trait datasets for each set of parameters with this method, using either a single tree specified (the species tree) or multiple trees specified (the gene trees).

As expected, we found that increasing the level of discordance results in an increasingly upward bias in estimates of the evolutionary rate from the species tree (Figure 4, green lines). As there are no internal branches in the species tree that can explain the increased trait covariances between non-sister taxa, such methods must propose a higher evolutionary rate to explain the data. In contrast, we found that both the covariance matrix approach (i.e. ***C\****; Figure 4A) and pruning algorithm approach (Figure 4B) yielded more accurate evolutionary rate estimates, ones that closely tracked the true population-scaled evolutionary rate as the level of discordance increased. Both phylogenomic comparative approaches can model the increased covariances generated by the increasing frequencies of discordant gene trees. As can be observed, both approaches tend to slightly underestimate the true evolutionary rate, but they are much closer to the true value than standard species tree estimates, especially at higher rates of discordance.

**Figure 4:**
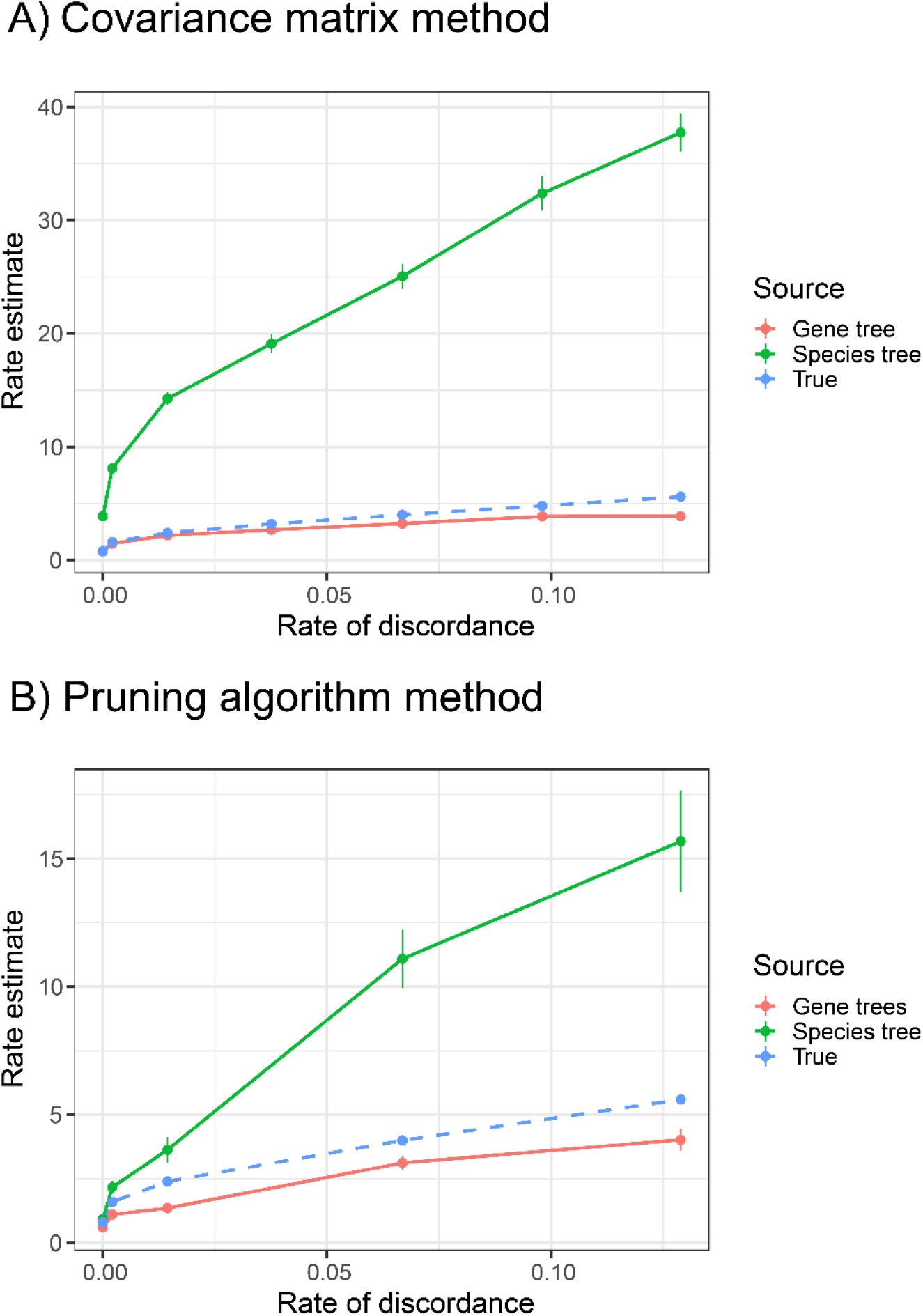
Phylogenomic comparative methods produce more accurate evolutionary rate estimates. A) Rate estimates obtained using a maximum-likelihood estimator applied to the covariance matrix (equation 5). B) Rate estimates obtained using numerical optimization of the likelihood with the pruning algorithm. In both panels, the green line shows inferences from methods using only the species tree, the red line shows the inferences from methods accounting for gene tree discordance, and the blue dashed line shows the true simulated value of the evolutionary rate. The level of gene tree discordance expected from each simulated species tree (see Supplementary Methods) is shown on the x-axis.

### Phylogenomic comparative approaches are robust to the effects of gene tree estimation error

In empirical datasets, it is reasonable to expect gene trees to be estimated with some degree of error, especially in the limits of short sequence length (such as ultra-conserved elements), long periods of evolutionary divergence, or high rates of sequencing error. In general, these sources of technical error should not be biased towards specific lineages, so their effect should be to cause general overestimation of gene tree discordance. This may in turn result in lower evolutionary rate estimates when using our approaches, as they might “overcorrect” the problem. More generally, we were concerned that increasing the rate of discordance might always leads to a lower evolutionary rate estimate, regardless of the true history that generated the data. Such behavior would present a potential problem for the application of our approaches to empirical datasets.

To address these concerns, we simulated traits from gene trees under a single, fixed rate of gene tree discordance (of approximately 15%) using the methods described in the previous section. We then applied our approaches to estimating σ^2^ to this dataset, varying the specified rate of gene tree discordance from 0 (in which case we used the standard species tree inference) to approximately 60%. In contrast to our initial concerns, we found that in both the covariance matrix (Figure 5A) and pruning algorithm (Figure 5B) approaches: 1) the effect of mis-specifying the rate of gene tree discordance is relatively small compared to the effect of using the species tree in place of gene trees; 2) increasing the specified rate of gene tree discordance leads to a small increase, rather than decrease, in the estimated evolutionary rate, but still closely tracked the true value. This latter effect may occur because increasing the specified rate of gene tree discordance requires branch lengths to be scaled down in accordance with *N*, resulting in less proposed time over which evolutionary changes can occur. Overall, these results suggest that gene tree estimation error should not be a major concern for our approaches, as long as the correct set of tree topologies is specified.

**Figure 5:**
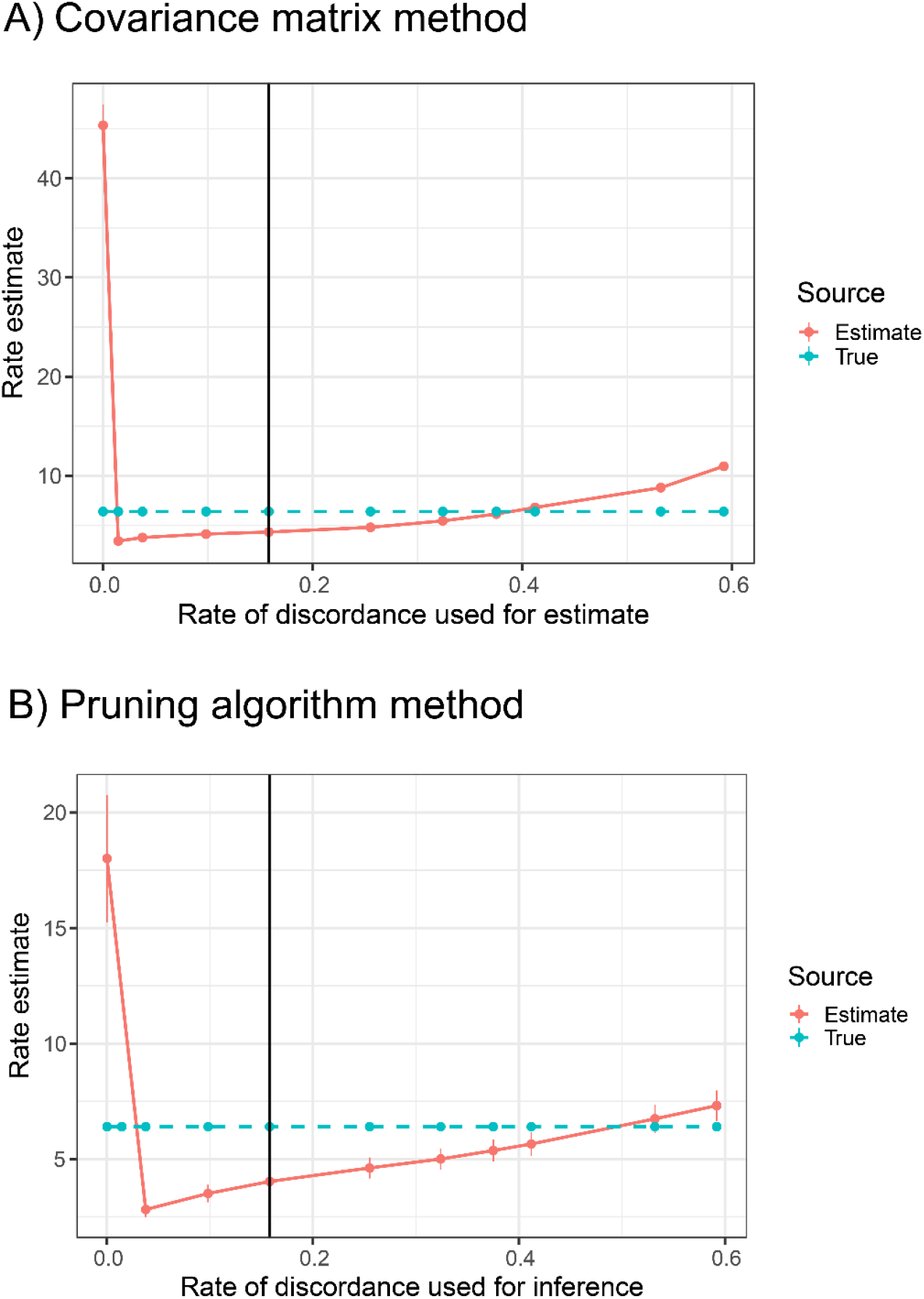
Phylogenomic comparative methods are robust to gene tree estimation error. In both panels, the solid vertical line denotes the true rate of discordance used to simulate the trait data, and the horizontal blue line denotes the true evolutionary rate. The x-axis shows the rate of discordance supplied to each approach when estimating the evolutionary rate from the simulated data. For a rate of discordance equal to 0, we used the standard species tree inference rather than gene tree inference. Specifying too much discordance can also cause overestimation of the evolutionary rate.

### Rate estimates for floral traits in the wild tomato clade *Solanum* are consistent with evolution on discordant gene trees

Our simulations show that when traits evolve on discordant gene trees, standard species tree approaches tend to greatly overestimate the true value, while our gene tree approaches slightly underestimate the true value but are much more accurate. The degree of discrepancy between species tree and gene tree approaches grows larger as the rate of discordance increases. To test further these expectations and to highlight the application of our methods to empirical data, we estimated the evolutionary rates of several floral traits (anther length, corolla diameter, and stigma length) measured in wild tomatoes (*Solanum*) (Haak et al. 2014). We obtained the time-scaled phylogeny of this clade from Pease et al. (2016) and converted from time in years to coalescent units assuming *N* = 100,000 and one generation every two years (Hamlin et al. 2020). We then pruned the phylogenetic tree into high and low ILS triplets, each consisting of three taxa. The high ILS group consisted of the following accessions (IDs from the Tomato Genetics Resource Center): *S. galapagense* LA0436, *S. cheesmaniae* LA3124, and *S. pimpinellifolium* LA1269; the low ILS group consisted of *S. pennellii* LA3778, *S. pennellii* LA0716, and *S. pimpinellifolium* LA1589. Based on the internal branch lengths in coalescent units, the high ILS and low ILS triplets had expected rates of discordance of approximately 47% and 0.9%, respectively. These rates correspond to the rates of discordance seen in empirically estimated gene trees in Pease et al. (2016). Based on our simulation results, if discordant gene trees contribute to tomato floral trait variation, we should see a greater discrepancy between species tree methods and our gene tree methods in the high ILS triplet.

In both of our approaches, we used the multispecies coalescent model to calculate the expected gene tree frequencies and branch lengths in each triplet. For the covariance matrix method, we used these expectations to construct the covariance matrix C* for each triplet using the *get_full_matrix*() method. For the pruning algorithm method, we specified a representative gene tree of each of the possible topologies with expected branch lengths, and weighted each tree by its expected frequency. For both methods, we used the standard species tree approach and our gene tree approaches to estimate the evolutionary rate using the mean trait values within each accession if multiple individuals were measured.

In line with our expectations, rate estimates obtained from standard species tree approaches are much higher than those obtained from both of our gene tree methods in the high ILS triplet, for all three traits (Figure 6). The discrepancy is much smaller in the low ILS triplets (Figure 6). The bias due to discordance was very large for the estimates obtained from the covariance matrix method in *seastaR* (Figure 6A), where the species tree rate estimates were several orders of magnitude higher than the gene tree estimates in the high ILS triplet. This is consistent with our simulation finding that the estimated evolutionary rate is more biased under discordance when using covariance methods (Figure 4A). Even when accounting for gene tree discordance, the rate estimates obtained from the covariance matrix method were substantially higher than those obtained from the pruning algorithm method (compare Figure 6A and 6B). This discrepancy can be explained by flat likelihood surfaces for the proposed values of σ^2^ (Supplementary Figure 1): the pruning algorithm method, which employs a likelihood search, reaches a likelihood plateau and does not propose further improvements, whereas the covariance matrix method uses the analytical likelihood estimator to obtain the maximum value, regardless of the shape of the likelihood surface. We believe the pruning algorithm estimates represent more biologically realistic rates of evolution.

**Figure 6:**
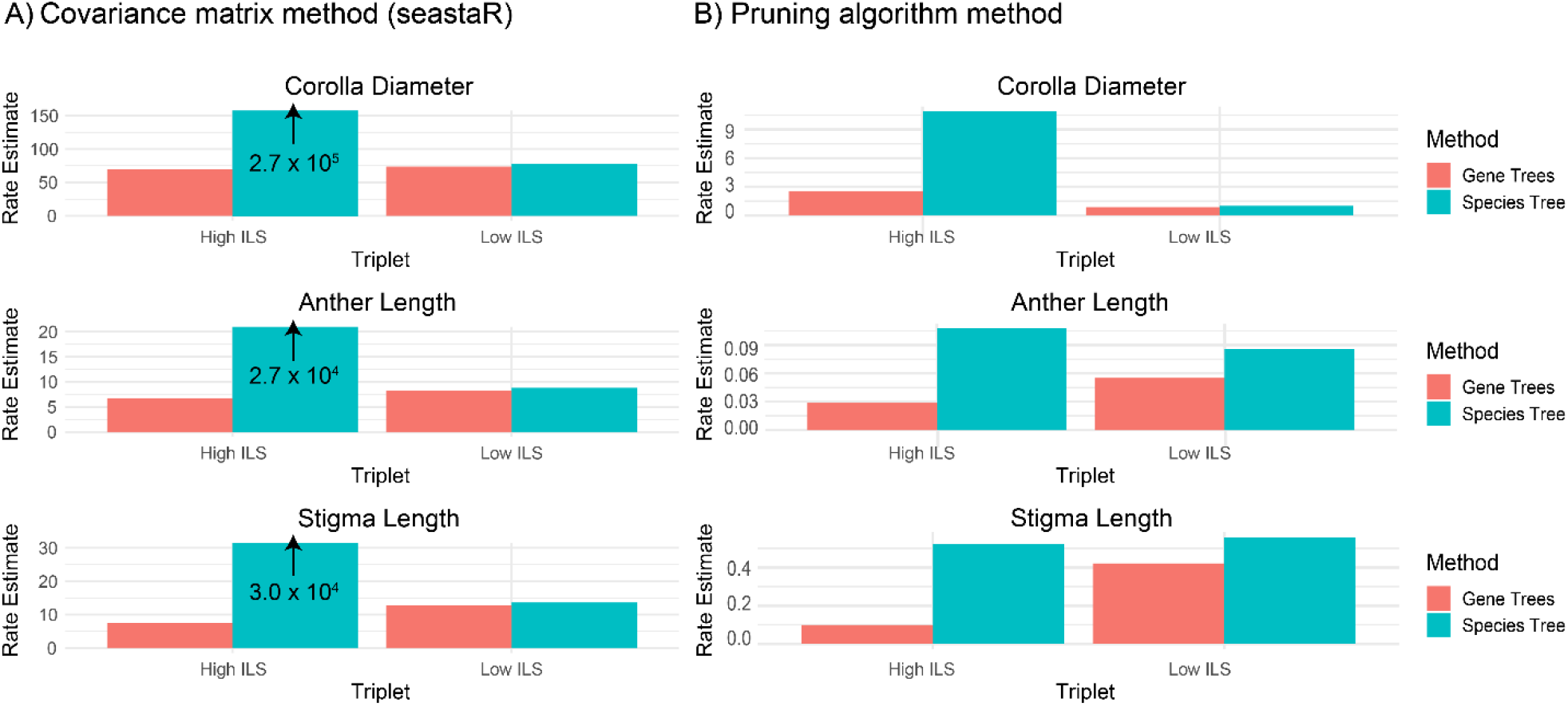
Evolutionary rate estimates for three floral traits in *Solanum* using our newly developed approaches (red bars), in comparison to standard species tree methods (blue bars). A) rate estimates (σ^2^) obtained using the analytical maximum-likelihood estimator as implemented in our R package *seastaR*. In the high ILS triplet, the species tree estimates (blue) go far above the scale of the y-axis, so these bars are labelled with an up arrow and the true estimated values for clarity. B) Rate estimates (σ^2^) obtained via maximum likelihood optimization using our pruning algorithm implementation for calculating likelihoods on gene trees. Note that the values on the y-axes are not the same in the two panels.

## Discussion

There has been much phylogenetic research focused on the accurate estimation of species trees in the face of gene tree discordance (e.g. Degnan and Rosenberg 2009; Bryant et al. 2012; Chifman and Kubatko 2014; Mirarab et al. 2014; Mendes and Hahn 2018; Zhang et al. 2018). Despite this focus on inferring trees in the face of discordance, standard phylogenetic comparative methods still rely on a single “resolved” tree to describe the shared history of species. Recent work has made it clear that, if only a single tree is used, gene tree discordance can shape trait variation and mislead comparative methods (e.g. Mendes et al. 2018; Hibbins et al. 2020). However, few solutions have been proposed to solve these problems, especially for quantitative traits evolving on clades containing discordance. Here, we have developed two approaches, which we refer to as *phylogenomic* comparative methods, that can incorporate gene tree discordance into comparative inference. One approach uses a more complete phylogenetic variance-covariance matrix that includes the covariance present in discordant gene trees. We have developed an R package, *seastaR*, for building this matrix using the frequencies and branch lengths of relevant gene trees. The second approach applies the pruning algorithm over a set of gene trees—concordant and discordant—to estimate likelihoods. Using simulation, we demonstrate that these methods generate more accurate evolutionary rate estimates for traits evolving in the presence of discordance, and are generally robust to the effects of gene tree estimation error. Finally, we demonstrate that empirical floral traits in the wild tomato clade *Solanum* are consistent with evolution on discordant gene trees, with the clade with a higher rate of gene tree discordance exhibiting a greater discrepancy in rate estimates between traditional approaches and our new methods.

Many phylogenetic comparative methods take the variance-covariance matrix, ***C***, as input (e.g. Pagel 1999; Housworth et al. 2004; O’Meara et al. 2006; Revell and Harmon 2008). Because of the wide use of ***C***, we anticipate that a more complete variance-covariance matrix, ***C\****, will be easy to incorporate into many comparative analyses. The *seastaR* package provides an easy way for users to generate ***C\****, either from a set of specified gene trees or from a specified species tree (assuming a multispecies coalescent process). Here, we have demonstrated how ***C\**** can be used to obtain a maximum-likelihood estimate of the rate of quantitative trait evolution under Brownian motion, a method that is also implemented in *seastaR*. One obvious extension of the use of ***C\**** is in phylogenetic generalized linear mixed models (PGLMMs), where the covariance matrix is often specified directly in packages such as *MCMCglmm* (Hadfield 2010). However, many popular packages for implementing comparative methods—such as *phytools* (Revell 2012), *ape* (Paradis and Schliep 2019), and *Geiger* (Pennell et al. 2014) —do not take a matrix directly, instead turning an input species tree into a matrix. Furthermore, they require a strictly bifurcating tree as input to construct a *phylo* class object. Integrating the ability to accept to take ***C\**** (or equivalent sets of gene trees) into these methods would enable a much larger array of inference tasks to take discordance into account.

The pruning algorithm is widely used in likelihood-based inference of phylogenetic trees (Felsenstein 1981) and for some applications in quantitative trait evolution (e.g. Hahn et al. 2005; FitzJohn 2012; Freckleton 2012; Ho and Ané 2014; Uyeda and Harmon 2014; Hiscott et al. 2016; Mitov et al. 2020). Our method using the pruning algorithm across a set of gene trees makes many of the same assumptions as previous implementations, but models each trait as the combined result of a large number of loci; these loci were represented in our calculations by a smaller number of exemplar gene tree topologies, each with the mean set of branch lengths for a given topology. Although it is not as straightforward to incorporate our method into other approaches as with ***C\****, because the pruning algorithm is a general method for calculating likelihoods, it has enormous potential to be applied to a wide variety of inference problems. As trees get larger, the computational cost of the matrix operations in equations 5 and 6 grows exponentially with the number of taxa. In contrast, the number of calculations in the pruning algorithm only grows linearly, and therefore trees with thousands of tips can be analyzed (Mitov and Stadler 2019). Furthermore, even though several methods for dealing with sparse matrices make it possible to analyze larger numbers of taxa (e.g. Hadfield and Nakagawa 2010), ***C\**** has more covariance entries and is therefore less sparse than ***C***; this again limits matrix-based approaches in phylogenomic comparative methods.

Both of our approaches can be extended in multiple ways. While we have only considered Brownian motion models here, there are multiple other trait models that could be used. The Ornstein-Uhlenbeck process is a popular model for trait evolution, with estimators available using both matrix (Hansen 1997; Butler and King 2004; Beaulieu et al. 2012; Rohlfs et al. 2014) and pruning algorithm approaches (FitzJohn 2012; Ho and Ané 2014; Uyeda and Harmon 2014; Mitov et al. 2020). Additional models for continuous traits include “early burst” (Harmon et al. 2010) and Lévy (“jump”) processes (Landis and Schraiber 2017). All of these models should be able to be accommodated by phylogenomic comparative methods. In addition, although we have described the covariances in our models with a particular set of gene trees in mind, both methods can be used with any weighted mixture of trees. This means that users do not have to assume a particular model of species tree evolution (e.g. the multispecies coalescent model) and can even ignore ILS altogether in favor of phylogenetic network models (e.g. Bastide et al. 2018).

There are also multiple caveats that come with our proposed approaches, and some important technical limitations to consider. First, errors in gene tree or species tree estimation might bias inferences. This is especially true if gene trees are being used as inputs, as we require both accurate and ultrametric trees. We found that error in gene tree frequencies and branch lengths is relatively inconsequential for our approaches, under the conditions considered here. Specifying a set of incorrect gene tree topologies may have more of an effect, but since ILS is expected to generate all possible topologies with respect to a single branch, we do not expect this to be a huge issue. Obtaining ultrametric gene trees remains challenging due to variation in rates of evolution among loci and small amounts of data per locus. Even when species trees are used to generate gene tree frequencies (i.e. *get_full_matrix*), many coalescent-based methods for inferring species trees do not estimate tip branch lengths (e.g. Liu et al. 2010; Mirarab et al. 2014), further limiting accurate inferences (but see Bastide et al. 2018; Hibbins et al. 2020). If there is uncertainty in the species tree topology or branch lengths, a straightforward solution would be to embed the approaches used here within a Bayesian framework (e.g. Huelsenbeck et al. 2000; Pagel and Meade 2006). It is important to note, however, that gene tree discordance is not equivalent to species tree uncertainty: averaging over each gene tree topology on its own in a Bayesian framework would simply mean averaging over many incorrect trees. Instead, a proper Bayesian approach to accommodating discordance would have to sum over a new set of gene trees (or covariances) for each species tree topology proposed, as was done here with a single topology.

A second caveat is that large numbers of taxa make it harder to accurately estimate both the matrix used in *seastaR* and the gene trees used within the pruning algorithm. If gene trees are predicted from theory, *seastaR* calculates ***C\**** from the species tree by breaking the tree into triplets. While this will return approximately correct covariances for all pairs of species, it necessarily ignores any covariance structures that might only be possible in trees with four or more taxa. The problem for the pruning algorithm approach could be even worse, as separate gene tree topologies must be specified: specifying representative gene trees for all possible topologies becomes prohibitive with more taxa because the number of gene trees grows super-exponentially. Even if gene trees are estimated from the data, with only a few dozen taxa there are more possible gene tree topologies than independent loci in a genome. Two solutions suggest ways around these issues. First, the problems can be somewhat ameliorated by recognizing that it is not the number of taxa that is the issue, but instead the number of lineages within “knots” (cf. Ané et al. 2007) on the larger phylogeny that are prone to gene tree discordance. For instance, even in a tree with 100 species, if only three are undergoing ILS, then only three topologies must be considered. Judicious choices as to the number of different topologies that must be considered in any particular analysis could save a lot of computational effort. Second, as mentioned in the Methods, one approach that can be applied to the pruning algorithm method is to sample a limited number of individual gene trees, either directly from the inferred trees or from the multispecies coalescent model applied to the species tree. Even if we have to sample 100 trees, the likelihood calculations on each are relatively fast and can be parallelized. Such a sampling scheme will also naturally recapitulate the degree of discordance associated with every branch in the species tree.

Throughout our analyses, we found that rate estimates using a single species tree differed from those accounting for gene trees, even when the level of discordance was very low or zero (Figure 4A). This result occurs because the two modes of inference are fundamentally different: even with no discordance, “gene tree” analyses are based on gene tree branch lengths, not species tree branch lengths. Gene tree branch lengths are always longer than species tree branch lengths because each pair of lineages is expected to coalesce 2*N* generations before their time of speciation (Gillespie and Langley 1979; Edwards and Beerli 2000). These longer gene tree branch lengths result in higher trait variances in the traits, such that a higher evolutionary rate must be proposed to explain the same data when using the species tree for analysis. This distinction highlights an additional challenge for a potential application of our pruning algorithm approach – ancestral state reconstruction. Because the internal nodes of gene trees—including concordant gene trees—do not exist at the same moment of time as the internal nodes of species trees, reconstructing ancestral states at the time of speciation requires knowledge of the contribution of each gene tree branch to trait evolution at that particular time point. This could be accomplished by the insertion of single-descendant nodes on gene tree lineages that are concurrent with ancestral nodes on the species tree. Inferring lineage-specific rate shifts will likewise require that each gene tree branch, or segment of a gene tree branch, be assigned to specific species tree lineages (cf. Ogilvie et al. 2017). In general, these considerations highlight the fact that using gene trees in place of a species tree is a fundamentally different mode of inference, and that standard comparative methods using the species tree may yield incorrect inferences even if there is no discordance.

In our analysis of flower morphological traits in wild tomatoes, we found that across both methods species tree rate estimates were much greater than gene tree estimates (Figure 6). For the high ILS triplet there was even more bias than in the low ILS knot. These results are consistent with a contribution of gene trees, rather than a single species phylogeny, to variation in these traits. Our analysis of gene tree error suggests that this result is not simply an artifact of increasing the specified rate of gene tree discordance, but is the result of biological variation in the floral traits. Furthermore, our findings have implications for the study of evolutionary rate variation among clades. For example, imagine that researchers wished to investigate whether the evolutionary rate of corolla diameter differed between our high ILS and low ILS triplets. Applying standard species tree methods, they would find that the corolla diameter of the high ILS species evolves at a much faster rate than in their low ILS counterparts. However, from our results in Figure 6, after correcting for the contribution of discordant gene trees, this difference disappears and the trait appears to evolve at approximately the same rate in both clades. This result highlights how variation in the rate of gene tree discordance among clades is a confounding factor when studying the evolution of lineage-specific rate shifts.

An increasingly common finding in phylogenomics is that of rapid and/or highly parallel trait evolution associated with rapid species radiations (Schluter et al. 1997; Boughman et al. 2005; Sun et al. 2012; Parins-Fukuchi et al. 2021; Urban et al. 2022). The application of classic comparative approaches in these systems has suggested that many radiations violate constant-rate Brownian motion models, with more complex models being proposed instead (Simpson 1944; Blomberg et al. 2003; Freckleton and Harvey 2006). However, adaptive radiations often have very little time between speciation events, resulting in high rates of gene tree discordance and therefore high potential for hemiplasy (Pease et al. 2016). Here we have found that apparently higher rates of trait evolution in rapid radiations may be perfectly consistent with a standard Brownian motion model with a constant evolutionary rate. In this circumstance, higher apparent rates of evolution are simply the result of a stronger contribution of discordant gene trees to covariance among species. Our proposed phylogenomic comparative methods help to address these issues, providing more accurate evolutionary inferences in systems with high rates of discordance.

## Acknowledgements

We would like to thank Ben Fulton and Jason Bertram for offering helpful programming and statistical advice in implementing the pruning algorithm approach, as well as David Haak for sharing floral trait data. This work was supported by the EEB Postdoctoral Fellowship awarded to Mark Hibbins by the Department of Ecology and Evolutionary Biology at the University of Toronto, as well as National Science Foundation grant DEB-1936187 awarded to Matthew Hahn.

