## Supplemental Materials and Methods for "Phylogenomic comparative methods: accurate evolutionary inferences in the presence of gene tree discordance"

### Supplementary Materials and Methods

### Parameters used for simulation study

As described in the main text, we used the model of Mendes et al. (2018) and varied the effective population size, $N$, to simulate trait evolution on gene trees with varying rates of discordance. In all simulations we used a three-taxon species tree with the topology ((A,B),C). We denote $t_1$ as the time of speciation of A a B, and $t_2$ as the time of speciation of C from the ancestor of A and B. In all simulations, we set $t_1$ = 4000 generations and $t_2$ = 50000 generations. We also used a constant value of $\mu\sigma_M^2$ = 0.0002. Each species tree has a single effective population size, $N$, which is shared across the whole tree. The branch lengths of this tree, which determine the rate of discordance and gene tree branch lengths, are obtained by dividing $t_1$ and $t_2$ by $2N$.

We modelled evolution along four gene trees: one concordant lineage-sorting tree with a frequency of $1 - e^{-\left(\frac{t_2}{2N} - \frac{t_1}{2N}\right)}$, and three (one concordant, two discordant) ILS trees each with a frequency of $\frac{1}{3}e^{-\left(\frac{t_2}{2N} - \frac{t_1}{2N}\right)}$. We only considered ILS as the source of discordance here, for the sake of simplicity. For each simulation condition, we compared standard inference using the species tree to our updated inferences that use information from gene trees. For each species tree we constructed the gene variance/covariance matrix, ***C****, using the expectations derived in Mendes et al. (2018). The covariance expectations are obtained by weighting the expected internal branch length of each gene tree topology by that topology's expected frequency, while the variance expectations are obtained by weighting the total heights of the gene trees. This gives the following expected variances and covariances:

$$Cov(AB) = 2N\mu\sigma_M^2 \left[ \left(1 - e^{-\left(\frac{t_2 - t_1}{2N}\right)}\right) \left( \frac{e^{\frac{t_2}{2N}} \left(\frac{t_2 - t_1}{2N}\right)}{e^{\frac{t_2}{2N}} - e^{\frac{t_1}{2N}}} \right) + \left(\frac{1}{3} e^{-\left(\frac{t_2 - t_1}{2N}\right)}\right) \right] \quad [1]$$

$$Cov(BC) = Cov(AC) = 2N\mu\sigma_M^2 \left(\frac{1}{3} e^{-\left(\frac{t_2 - t_1}{2N}\right)}\right) \quad [2]$$

$$Var(A) = Var(B) = Var(C) =$$

$$2N\mu\sigma_M^2 \left[ \left(1 - e^{-\left(\frac{t_2 - t_1}{2N}\right)}\right) \left(\frac{t_2}{2N} + 1\right) + \left(e^{-\left(\frac{t_2 - t_1}{2N}\right)}\right) \left(\frac{t_2}{2N} + 1 + 1/3\right) \right] \quad [3]$$

We simulated trait values by using the R function *mvrnorm* to make random draws from a multivariate normal distribution specifying ***C**** as the covariance structure. These data were used as the input to both methods of inference.

For inferences based on the estimator of $\sigma^2$ using a matrix, we passed $\boldsymbol{C}$ and $\boldsymbol{C^*}$ to Equation 5 of the main text for species tree and gene tree inferences, respectively. For inferences based on the pruning algorithm, we passed our software the species tree in coalescent units for the species tree inference mode. For the gene tree inference mode, we calculated the expected branch lengths within each of the four gene trees using the multispecies coalescent (Hibbins and Hahn 2019) and gave these representative gene trees to the software along with their expected frequencies. For the concordant lineage-sorting tree, these expected branch lengths are:

$$E[t_{A-B}] = \frac{t_1}{2N} + \left(1 - \frac{\frac{t_2 - t_1}{2N}}{e^{\left(\frac{t_2 - t_1}{2N}\right)} - 1}\right) \qquad [4]$$

for the timing of the first coalescent event (going backwards in time), and

$$E[t_{B-C}] = E[t_{A-C}] = \frac{t_2}{2N} + 1 \qquad [5]$$

for the timing of the second coalescent event. The internal branch for this gene tree is calculated as Equation S5 minus Equation S4. For the three gene trees arising from ILS, we have

$$E\left[t_{first}\right] = \frac{t_2}{2N} + 1/3 \qquad [6]$$

for the first coalescent event, and

$$E[t_{second}] = \frac{t_2}{2N} + 1/3 + 1 \qquad [7]$$

for the second.

For the results presented in Figure 4, when applying the matrix-based estimator (Figure 4A), we simulated data using specified values of *N* of 2000, 4000, 6000, 8000, 10000, 12000, and 14000, corresponding to levels of gene tree discordance of 0.00067%, 0.212%, 1.44%, 3.76%, 6.68%, 9.8%, and 12.8%, respectively. For each condition we ran 1000 simulations. When applying the pruning algorithm method (Figure 5.4B), we simulated data using specified values of *N* of 2000, 4000, 6000, 10000, and 14000, corresponding to levels of gene tree discordance of 0.00067%, 0.212%, 1.44%, 6.68%, 9.8%, and 12.8%, respectively. For each of these conditions we ran 100 simulations.

For the results presented in Figure 5, we simulated the data using *N* = 16000, corresponding to a rate of discordance of 15.8%. For both the matrix-based and pruning algorithm estimators, we specified values for the inference of *N* = 2000, 6000, 8000, 12000, 16000, 24000, 32000, 40000, 48000, 80000, and 150000, corresponding to rates of discordance of 0.00067%, 1.44%, 3.76%, 9.8%, 15.8%, 25.5%, 32.4%, 37.5%, 41.2%, 53.2%, and 59.2%. We ran 1000 simulations per condition for the matrix-based approach, and 100 simulations per condition for the pruning algorithm approach.

**Calculation of likelihood surfaces for tomato trait data**

In our analysis of tomato floral traits, we found that the covariance matrix method generally returned higher evolutionary rate estimates than the pruning algorithm method (Figure 6, main text). We suspected that this was because of the presence of flat likelihood surfaces for our traits. To verify this, we used the analytical likelihood formula for a given value of $\sigma^2$ as reported in O'Meara et al. (2006). We used this formula to calculate the likelihood for 400 evenly spaced values of $\sigma^2$ between 0 and 200, for each trait and within both phylogenetic triplets. As expected, our traits had flat or gently sloping likelihood surfaces, which would have prevented the pruning algorithm likelihood optimization from climbing the surface to the analytical maximum likelihood value (Supplementary Figure 1).

### Supplementary Figures

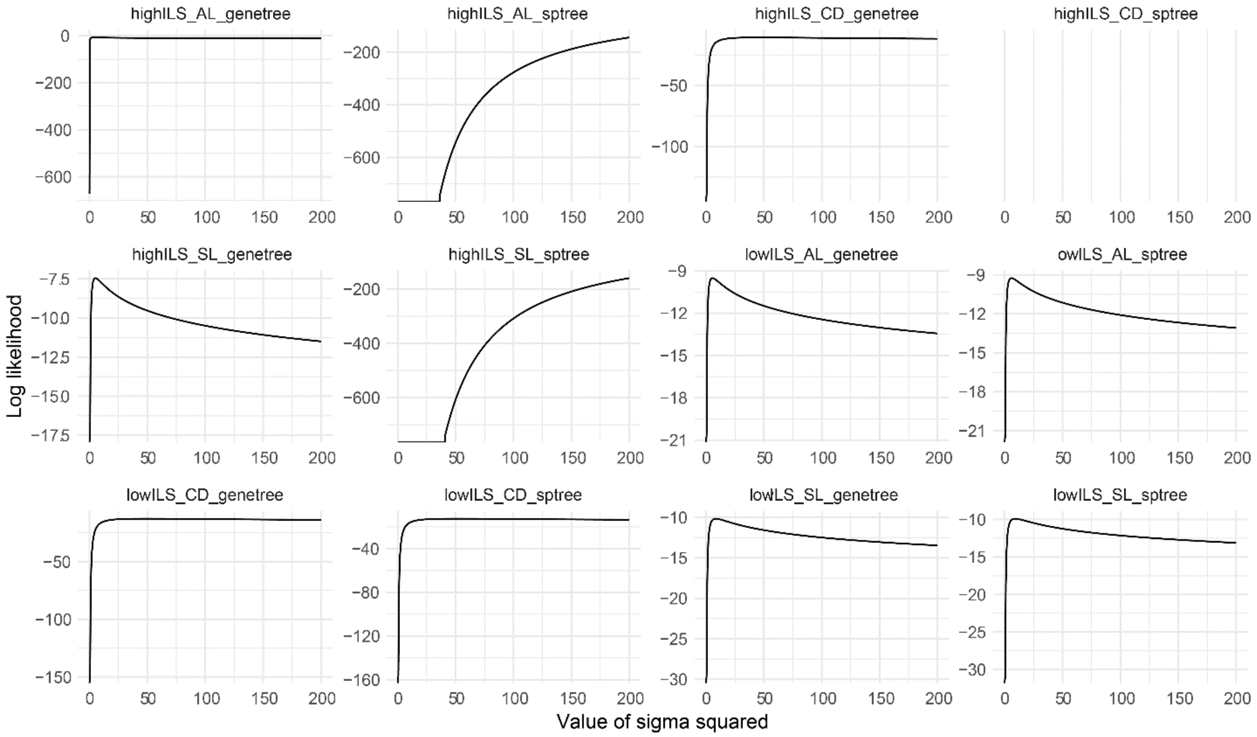


*Supplementary Figure 1*: Likelihood surfaces, calculated using the analytical estimator in Equation 3 of O'Meara et al. (2006), for each analysis of the empirical tomato trait dataset. In the individual plot captions, AL, CD, and SL refer to anther length, corolla diameter, and stigma length respectively. In "sptree" plots we supplied the standard covariance matrix ***C*** to the likelihood calculation, and in "genetree" plots we used ***C****.
